# Abiotic factors are the primary determinants of endemic Hawaiian *Drosophila* microbiome assembly

**DOI:** 10.1101/2025.05.06.652154

**Authors:** Matthew J. Medeiros, Sean Schoville, Donald Price, Joanne Y. Yew

## Abstract

The Hawaiian *Drosophila* radiation exemplifies rapid adaptation and species diversification. Many factors have been attributed to these phenomena, including allopatry, sexual selection, and ecological specialization. In recent years, the microbiome has come to the forefront as an important driver of adaptation that is capable of facilitating host survivorship, enhancing resilience to local environmental challenges, and enabling the use of different dietary resources. To determine how microbial communities assemble in natural populations and potentially contribute to the rapid adaptation of Hawaiian drosophilids, we conducted a survey of bacterial and fungal communities from over 500 wild flies collected from across six islands of the Hawaiian archipelago. These samples represent a breadth of host plant specializations, habitats, lifestyles, and endemicity. Our findings reveal that microbiome assembly is largely driven by abiotic factors including elevation, temperature, rainfall, and evapotranspiration, but is not strongly constrained by phylogenetic relatedness. Identical species inhabiting two separate locations exhibited different microbiomes. By contrast, distantly related species inhabiting the same site had more similar microbiomes. The microbiomes of native species also differ from recently introduced, non-native *Drosophila* in terms of diversity, composition, and function. Given the myriad roles of the microbiome in nutrition, reproduction, and mate choice, these results support a role for the microbiome in the remarkable ecological divergence of Hawaiian *Drosophila*.

## INTRODUCTION

Animal evolution takes place in a natural world teeming with pathogenic and beneficial microorganisms, both of which shape a breadth of physiological functions including digestion, detoxification, immune response, circadian rhythm regulation, thermal tolerance, and reproduction (Baldassarre et al 2022, Kohl et al 2022, Kucuk 2020, Lee and Hase 2014, Li et al 2021, Suyama et al 2023, Tefit et al 2023, Walters et al 2020, Zhang et al 2024). The complex partnership between host and microbe has been hypothesized to facilitate adaptation by improving resilience to climate stress, offering protection from pathogens and parasites, and enabling a move to novel dietary niches (Gao et al 2023, Kohl et al 2014, Zhang et al 2024). How host-associated microbiomes assemble and contribute to adaptation largely remains an open question. In humans and other animals, environmental factors, especially diet, significantly influence the composition and diversity of the microbiome (Carmody et al 2015, Turnbaugh et al 2009). Additionally, organisms may co-evolve with microbial taxa over evolutionarily long periods of time, referred to as phylosymbiosis (Brooks et al 2016), and implement genetic controls that regulate bacterial diversity and/or abundance (Chaston et al 2016). Neotropical butterflies are one example of this phenomenon whereby host phylogeny plays a prominent role in predicting gut microbial communities (Ravenscraft et al 2019). Other genera exhibit different relative contributions of habitat versus phylosymbiosis in host microbiome assembly (e.g., (Lim and Bordenstein 2020, Martoni et al 2023, Perez-Lamarque et al 2022).

The extreme geographic isolation of Hawai□i as well as its diverse climate conditions contribute to the high endemism and biodiversity of plants and animals found in the archipelago (Barton et al 2021). The rapid adaptive radiation of the Hawaiian *Drosophila* provides an excellent opportunity to investigate the relative contributions of endemic status, genetics, evolutionary history and environmental factors to the assembly of microbial communities. The bulk of *Drosophila* diversification occurred in the Hawaiian Islands, where a recent radiation (∼10 Ma; (Church and Extavour 2022, Magnacca and Price 2015) has resulted in hundreds of species across a number of major clades, almost all of which are single-island endemics. The flies exhibit a variety of dietary habits: the larvae of some species are dietary generalists and subsist on several different host-plants, while others are strict specialists and associate with a single plant clade (Magnacca et al 2008, Magnacca and Price 2012, Magnacca and Price 2015). The Hawaiian Islands also offer a variety of abiotic environmental conditions along a chain of islands of known age (Clague 1996). This “natural laboratory” allows island biogeographic hypotheses to be tested with respect to host-associated microbiomes, such as the prediction that taxonomic diversity increases with island age (Hembry et al 2021, Matthews et al 2021, Shaw and Gillespie 2016). Other geographical aspects including latitude, elevation, and area have also been shown to shape macroorganism diversity (Davison et al 2018, Grayson and Lennstron 2022, Losos and Ricklefs 2009) but it remains unclear whether these patterns are recapitulated by host-associated microbiomes (Brown et al 2023, Dickey et al 2021, Fierer and Jackson 2006, Meyer et al 2018).

Here we describe the bacterial and fungal profiles of wild native and non-native *Drosophila* in order to understand the influence of environmental conditions, lifestyle, and evolutionary history to microbiome assembly. We performed high throughput amplicon sequencing of the 16S rRNA gene and ITS region to characterize, respectively, bacterial and fungal communities from over 500 endemic and invasive flies sampled from six islands of the Hawaiian archipelago and looked for evidence of whether the following features influence microbial diversity: geographic location and climate, genetic relatedness, larval dietary habits, endemic status, and island age.

## METHODS

### Wild fly collection and processing

We sampled over 500 flies in the field from 20 locations across six islands (**Supp. Table 1**). A subset of 419 flies were used for 16S rRNA analysis and 510 flies for ITS analysis. The elevations of collection sites ranged from 599 – 1536 m, average annual rainfall ranged from 750 – 6392 mm, average annual temperature ranged from 13 - 19.8 °C, and average annual evapotranspiration ranged from 599 -978 mm (**Fig. 1**).

**Figure 1.**
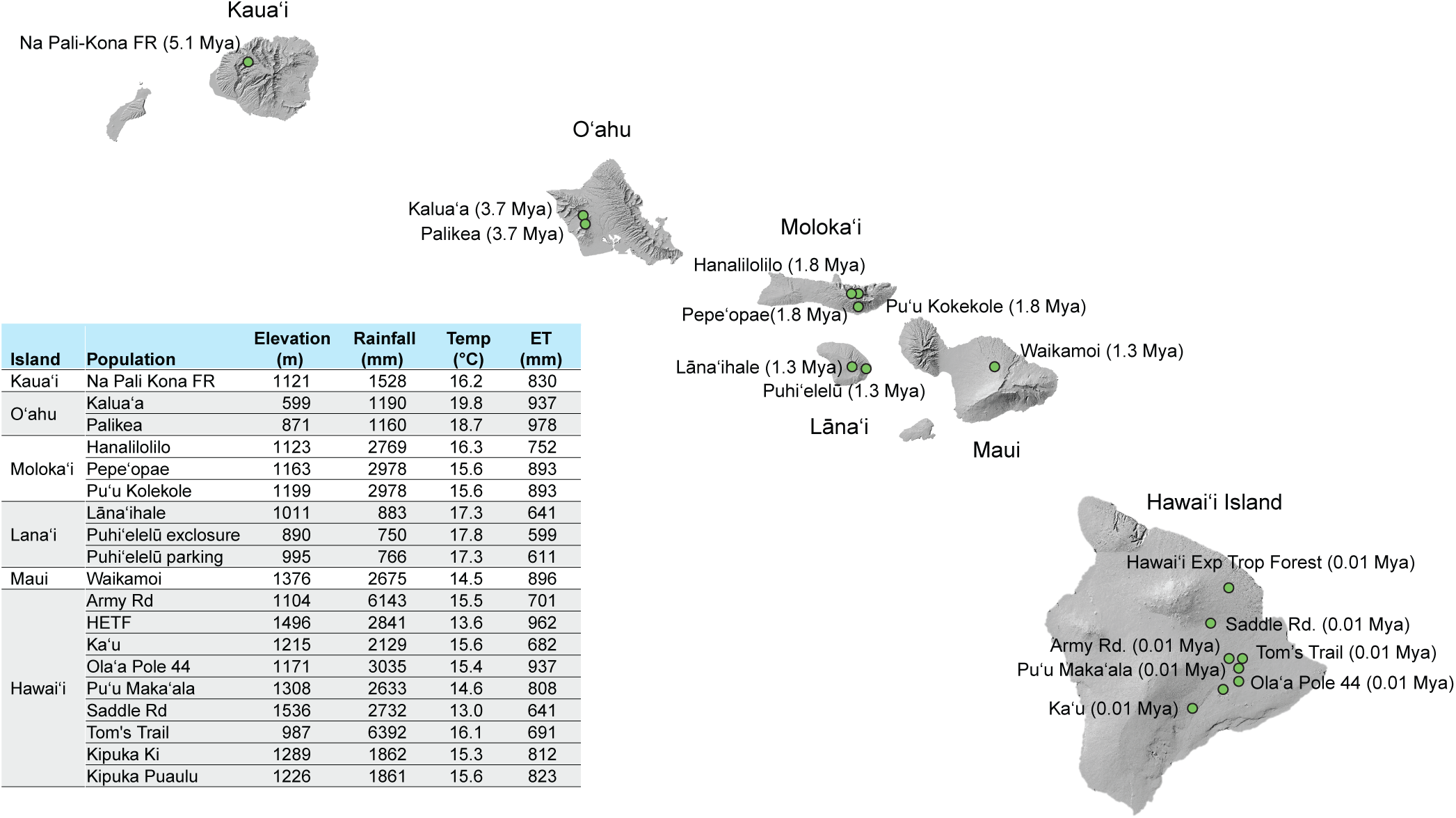
*Drosophila* collection sites, corresponding volcano ages, and environmental conditions. Environmental conditions are based on Giambelluca et al. (2013) and represent average annual values. Volcanic age estimates are based on Carson and Clague (1995); Mya: million years ago; HETF: Hawai‘i Experimental Tropical Forest; ET: evapotranspiration.

Wild flies were captured on sponges baited with crushed Cavendish bananas (*Musa acuminata*) mixed with baker’s yeast at a ratio of approximately 5 g of yeast per banana as well as fermented mushroom (*Agaricus bisporus*) “tea.” The mushroom tea was made by combining 200 g mushrooms in 1 L of water and allowing the mixture to ferment at room temperature for 14 d. Live flies that landed on the bait were captured with a polypropylene tube and transferred to 95% EtOH within 10 min and stored in coolers with ice packs. Samples were transferred to - 10 °C storage within 1-2 hrs of collection.

### Genomic DNA extraction and library preparation

DNA was extracted using published procedures (Medeiros et al 2024). Briefly, flies were surface sterilized with 2 washes in 95% EtOH followed by 2 washes in sterile water. Individual flies were homogenized in ATL buffer from PowerMag Bead Solution (Qiagen; MD, USA) with 1.4 mm ceramic beads (Qiagen) using a bead mill homogenizer (Bead Ruptor Elite, Omni, Inc; GA, USA) and extended vortexing for 45 min at 4 °C. The homogenate was treated overnight with proteinase K (2 mg/ mL) at 56 °C and DNA was extracted using the MagAttract PowerSoil DNA EP Kit (Qiagen) according to manufacturer’s instructions. Bacterial diversity was characterized by PCR amplification of the 16S rRNA gene with primers to the V3-V4 region (515F: GTGYCAGCMGCCGCGGTAA; 806R: GGACTACNVGGGTWTCTAAT) (Parada et al 2016). Fungal diversity was characterized using primers to the internal transcribed spacer (ITS1f: CTTGGTCATTTAGAGGAAGTAA; ITS2: GCTGCGTTCTTCATCGATGC) (White et al 1990). The primers contain a 12-base pair Golay-indexed code for demultiplexing.

The PCRs were performed with the KAPA3G Plant kit (Sigma Aldrich, MO, USA) using the following conditions: 95 °C for 3 min, followed by 35 cycles of 95 °C for 20 seconds, 50 °C for 15 seconds, 72 °C for 30 seconds, and a final extension for 72 °C for 3 min. The PCR products were cleaned and normalized with the Just-a-plate kit (Charm Biotech, MO, USA). High throughput sequencing (HTS) was performed with Illumina MiSeq and 250 bp paired-end kits (Illumina, Inc., CA, USA). Each individual *Drosophila* collected was identified using both morphology and a Genbank BLAST search of the approx. 800 base COI barcoding region.

### Microbiome data analysis

Post-processing of HTS data (read filtering, denoising, and merging) was performed using the “MetaFlow|mics’’ microbial 16S pipeline for bacteria and the fungal ITS pipeline for fungi (Arisdakessian et al 2020). Reads shorter than 20 bp and samples with fewer than 5,000 reads were discarded. Paired reads are merged if the overlap is at least 20 bp with a maximum 1 bp mismatch. The contigs generated by DADA2 (Callahan et al 2016) were processed using MOTHUR (Schloss et al 2009) and initially aligned and annotated using the SILVA v138 database (Quast et al 2013). For all analyses, we chose a 97% sequence similarity cutoff for determination of OTUs as we were attempting to investigate broad patterns rather than investigate the effects of specific strains of microbes. When numerically abundant sequences were unassigned by the pipeline, we identified them by manual searches in NCBI BLAST, UNITE (Nilsson et al 2019), and MycoBank (Robert et al 2013) using a >95% sequence similarity cutoff. The ITS data were normalized in R by transforming sample counts to proportions (McKnight et al 2019). Analyses were performed after clustering at the genus level using the *phyloseq* package (McMurdie and Holmes 2013) in R version 4.3.1 (RCoreTeam 2022). Relative abundance charts were constructed after grouping flies from the same geographical area or taxon. Univariate multiple testing with an F-test was used to test for significant differences in microbial taxa (*phyloseq*). Because we observed no difference in the microbiomes of female and male flies, these samples were treated as independent replicates unless otherwise noted.

We generated several datasets to test focal questions, including a subset that includes only endemic Hawaiian *Drosophila* (“HD”), and a subset of flies introduced to Hawai□i (“nonHD”). For the HD and nonHD datasets, we first examined whether the dispersion (multivariate variance) in microbial community composition was unequal among fly species. We employed the function *betadisper* in the *vegan* package (Oksanen et al 2015) to test for heterogeneity of multivariate dispersions, using the *anova* function in the *R stats package* to test for significance. We calculated dispersion among samples based on Bray-Curtis dissimilarity values (Bray and Curtis 1957), which is expected to perform robustly as the metric exhibits high sensitivity to group-level differences and linearity in abundance transforms across samples (Kers and Saccenti 2021, Ricotta and Podani 2017). We selected alternative downstream statistical tests depending on whether the among-group dispersion was homogeneous (*i.e.* permutational multivariate analysis of variance, PERMANOVA), or heterogeneous (*i.e*. Mantel correlation and multiple regression on distance matrices, MRM). Although Mantel tests are sensitive to heterogeneity in dispersions, we employ them as an initial omnibus test in conjunction with logistic regression using MRM, which is considered to be robust to among-group variance and an unbalanced sampling design (Anderson and Walsh 2013, Lichstein 2007, McArtor et al 2017, Warton et al 2012).

### Phylosymbiosis

To test for phylosymbiosis, a pattern where the microbial community would have greater similarity among closely related species, we employed Mantel tests to examine the association between microbial composition dissimilarity and phylogenetic dissimilarity. We included only endemic Hawaiian *Drosophila* samples to align with a recent Hawaiian drosophilid phylogeny based on phylogenomic data (Church and Extavour 2022). As several Hawaiian *Drosophila* are missing from the phylogeny, we placed them with known sister taxa as unresolved terminal branches: *D*. *grimshawi* was placed with *D. pullipes* and *D. craddockae*; *D. oreas* was placed as sister to *D. conspicua*. We calculated the pairwise phylogenetic distance among *Drosophila* taxa using branch lengths, through the *cophenetic.phylo* function in the package *ape* (Paradis and Schliep 2019). We used the *mantel* function in the *vegan* package to test for a correlation between the community dissimilarity and phylogenetic dissimilarity. A non-parametric Spearman’s rank correlation was used in the *mantel* function and the data were permuted 9,999 times to calculate significance of the correlation coefficient.

### Abiotic factors

To test for the role of abiotic factors in microbial community assembly, we first assessed a set of climatic predictors for independence using Pearson’s correlation, including temperature, elevation, rainfall, and evapotranspiration (Giambelluca et al 2013). As temperature and elevation were highly correlated, we focus on elevation as a predictor. We also assessed island age by assigning a value based on dates in Carson & Clague (Carson and Clague 1995), with Kaua’i being oldest, followed by O’ahu, Moloka’i, Lāna’i, Maui, and Hawai□i Island being the youngest. Island size typically correlates with island age, with younger islands being larger than older ones, but this pattern does not apply to Moloka’i and Lāna’i, which used to be joined with Maui as one larger island. Lāna’i was dropped from our analyses of island age and size due to collecting only two native *Drosophila* from that island.

### Island age and community diversity

To minimize potential bias from uneven sampling between each island, we randomly subsampled 45 individuals (for 16S rRNA comparison) and 56 individuals (for ITS comparison) from each island. Samples from Lāna’i were omitted since only two native flies were collected.

### Environment, geography, and phylogeny

To test the relative importance of phylogenetic relatedness, and environmental and geographical variables, we used multiple regression to test phylogenetic distance, elevation, evapotranspiration, rainfall, and island age as explanatory factors in microbial community composition. Each variable was analyzed using the MRM function in the *ecodist* package (Goslee and Urban 2007). We used the logistic method, as the among-group variance was heteroscedastic. A desirable feature of logistic regression is that it does not assume linear dependence between the dependent and independent variables, although it does assume that the independent variables are linearly related to the log odds. The effect size and statistical significance of each variable was calculated using 1,000 permutations of the data. The overall deviance of the logistic regression model can be used as a measure of the goodness of fit, with values closer to zero showing improved model performance. To help visualize the effect of each variable graphically, we employed distance-based redundancy analysis using the *capscale* function in the *vegan* package. We regressed a distance matrix of each predictor variable against the Bray-Curtis dissimilarity matrix of the microbes in a linear model, and used ordination to display the first two principal coordinates. Samples were visually grouped in the ordination according to their value of the predictor variable.

To further explore the predictive power of environmental variables on microbiome variation, we examined the effect of among-site variation within endemic species using PERMANOVA. PERMANOVA is a non-parametric multivariate statistical permutation test that employs distance matrices for response and predictor variables to test group-level differences. We restricted these analyses to two species with large sample sizes (>30 samples from at least 6 sites) to ensure sufficient degrees of freedom in the statistical tests: *D. tanythrix* (55 samples, 8 sites) and *D. sproati* (36 samples, 7 sites). We used the *adonis2* function in the *vegan* package to conduct the PERMANOVA (McArdle and Anderson 2001). We also used *D. sproati* and *D. tanythrix* to investigate whether gut microbial beta-diversity communities differed within or between sites.

### Dietary specialization

To test the impact of dietary specialization on microbiome composition, we examined a subset of picture-wing flies that designated as generalists (16S rRNA: n=30; ITS: n=42) and specialists (16s rRNA: n=184; ITS: n=190) based on field observations of larval association with host plants (Magnacca et al 2008). Generalists in our data set include *D. villospedis*, *D. crucigera*, and *D. grimshawi*. For specialists, we include *D. ochracea*, *D. sproati*, *D. prolaticilia*, *D. punalua*, *D. ambochila*, *D. inedita*, *D. hirtitibia*, *D. oreas*, *D. murphyi*, *D. cilifera*, *D. melanocephala*, *D. orphnoopeza*, *D. anomalies*, *D. quasianomalipes*, *D. picticornis*, *D. planitibia*, *D. neoperkinsi*, *D. bostrycha*, *D. limitata*, *D. setosimentum*, and *D. cilaticrus*.

### Functional analysis

In order to determine if bacterial microbiome functional differences varied among fly species, we used PICRUSt2 (Douglas et al 2020). PICRUSt uses the 16S rRNA marker gene to match microbial OTUs and their reference genomes, where the functional annotation of each microbial genome is retrieved. Functional annotations of gene families present in each genome are based on orthologs within the Kyoto Encyclopedia of Genes and Genomes (KEGG). We then used STAMP v2.13 (Parks et al 2014) to analyze differences among fly groups based on functional abundance and to generate visualizations of the functional data. The functional profiles were filtered to retain features based on statistical significance (*p =* 0.01) and remove features with few supporting reads (maximum per group < 1).

## RESULTS

### Differences between Invasive and endemic drosophilid species in Hawai***U***i

The Hawaiian *Drosophila* radiation spans the eight main islands of the Hawaiian archipelago and exhibits a range of diets, distinct host plant preferences, and habitats (Magnacca et al 2008). Non-native drosophilids including cosmopolitan species such as *D. suzukii* and *D. immigrans* are also commonly found throughout Hawai‘i. To determine whether the microbiome composition is specific to the evolutionary history of Hawaiian flies, we compared the bacterial and fungal compositions of 60 native species (including 29 from the picture-wing subgroup) and 6 introduced species. We hypothesized that the microbiomes of native flies will be i) distinct from introduced species in terms of composition and predicted function, ii) differentially affected by environmental factors, and iii) characteristic of host dietary specialization.

High –throughput amplicon sequencing of the bacterial marker 16S rRNA and fungal marker ITS revealed that the microbiomes of both native and non-native species contain similar numbers of bacterial genera (26 ± 6 in endemic species vs. 22 ± 7 in cosmopolitan species) and fungal genera (10 ± 4 in endemic species vs. 9 ± 3 in cosmopolitan species). Members of the *Enterobacteriaceae*, *Pseudomonas*, and *Dysgonomonas* genus were among the top 10 taxa in all flies regardless of endemic status (**Fig. 2**). With respect to the fungal microbiome (mycobiome), eight genera were shared between endemic and non-endemic flies (**Fig. 2**). The dominant class in both populations was the yeast *Saccharomycetes*, consistent with previous reports of Hawaiian drosophilids (O’Connor et al 2014) (**Supp. Table 2**). Despite the overlap in common bacterial and fungal taxa, Hawaiian *Drosophila* exhibit bacterial communities with greater taxonomic richness and evenness compared to non-native flies (**Supp. Fig. 1**) and were compositionally distinct from invasive species (**Fig 3A**). In particular, several pathogenic and commensal bacteria species differed in abundance with endemic status (**Supp. Table 2, Fig. 2B**). Functional analysis of endemic and invasive bacterial microbiomes using PICRUSt2 suggests slight shifts in functional space, rather than the gain or loss of functional pathways (**Fig 2C**).

**Figure 2.**
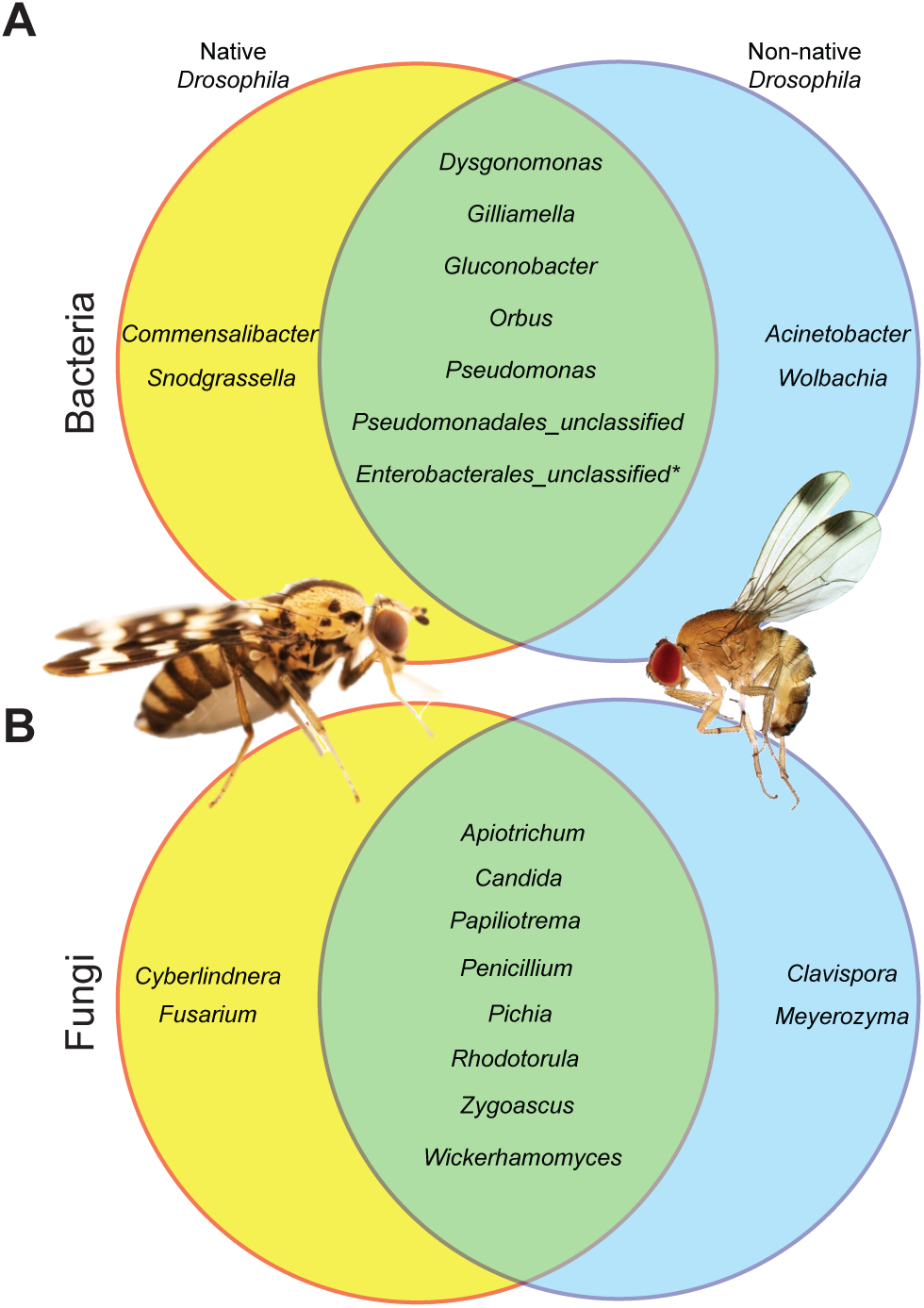
Venn diagrams of microbial taxa found in native and non-native drosophilids, based on the ten most abundant (**A**) bacterial and (**B**) fungal genera; *likely *Enterobacteriaceae* and *Yeriniaceae*. Photo credit: *D. grimshawi* (left; J. Sartore and Hawai‘i Invertebrate Program) and *D. suzukii* (right; V. Stacconi (Stacconi 2022)).

**Figure 3.**
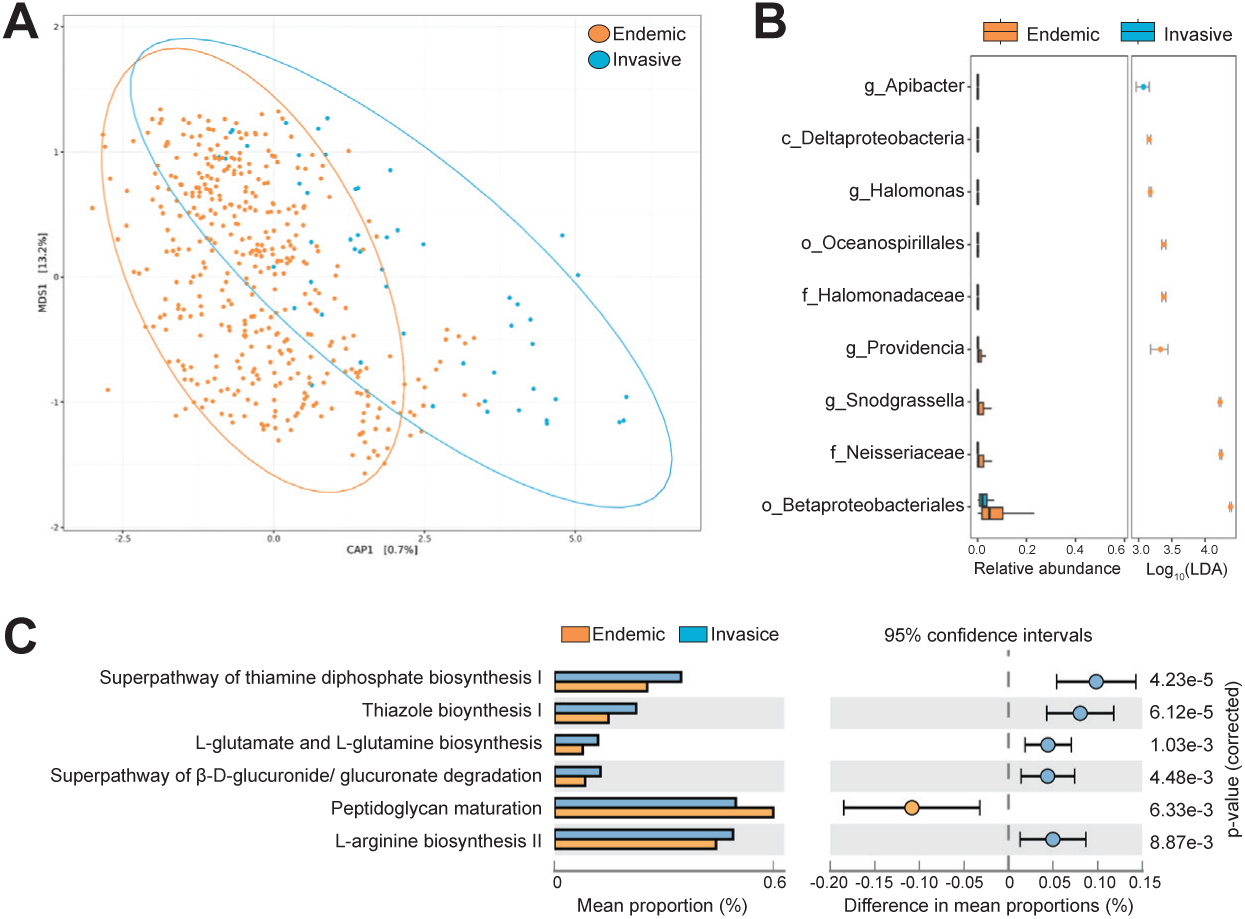
Visualization of differences in the composition and predicted function of bacterial communities in endemic Hawaiian *Drosophila* and non-native flies. (**A**) Ordination employing a distance-based redundancy analysis (dbRDA) of Bray-Curtis distance in relation to invasive species status. (**B**) Differential abundance analysis showing bacterial groups that differ significantly among the non-native (orange) and endemic fly species (blue). Significant differences at the lowest taxonomic level are shown. (g: genus; c: class; o: order; f: family). (**C**) Predicted functional differences between endemic Hawaiian *Drosophila* and non-native species. Functional profiles are based on KEGG annotations of microbial reference genomes and are weighted by bacterial abundance in each fly sample.

The shift in bacterial microbial diversity was associated with changes in its functions related to several notable biosynthetic pathways. Invasive species host microbes that are enriched for peptidoglycan maturation, which is important for maintenance of cellular integrity and enables survival in challenging environments. In contrast, endemic species host microbes that are enriched for biosynthesis of thiazole and thiamin (vitamin B), essential micronutrients, as well as L-glutamate and L-glutamine biosynthesis (ammonia assimilation), L-arginine biosynthesis (production of metabolites and amino acids), and beta-D-glucuronide and D-glucuronate degradation, both of which are related to detoxification processes.

To evaluate the effect of environmental variables on the microbiome composition of native and non-native fly species, we employed a multiple regression on distance matrices (MRM). Our analyses reveal that bacterial communities varied significantly with elevation, evapotranspiration, island age, and endemic status, with elevation showing the strongest effect on microbiome dissimilarity among samples (**Table 1; Supp. Fig 2**). As with bacterial communities, fungi of native species were affected by elevation, evapotranspiration, and rainfall (listed in order of explanatory strength) (**Table 2**). Notably, when the bacterial and fungal profiles of invasive species are examined separately for environment-induced variation using an MRM, no significant effects were found (**Supp. Table 3; Supp. Fig 2**).

**Table 1.** Multiple regression on distance matrices (MRM) analysis of native and non-native *Drosophila* bacterial profiles and environmental and genetic predictors. ^1^Microbiome distance values among samples are based on Bray-Curtis dissimilarity values, whereas predictor variables are measured as Euclidean distance values. All predictor variables are shown and significant predictors are in bold-face (residual deviance = 12220.36).

**Table 2.** Multiple regression on distance matrices (MRM) analysis of native and non-native *Drosophila* fungal profiles and environmental and genetic predictors. ^1^Mycobiome distance values among samples are based on Bray-Curtis dissimilarity values, whereas predictor variables are measured as Euclidean distance values. All predictor variables are shown and significant predictors are in bold-face (residual deviance = 28438.57).

### Testing for phylosymbiosis in Hawaiian Drosophila

Considering that endemic hosts harbor different microbes than invasive species, we next asked whether the microbiome of Hawaiian flies is linked to host phylogeny and evolutionary history. We looked for evidence of phylosymbiosis by testing whether more closely related host species will exhibit more similar microbial profiles based on Bray-Curtis distances. We found statistically significant, but weak relationships between bacterial and fungal diversity and phylogenetic relatedness (16S rRNA: non-parametric Mantel statistic *r* = 0.0727, *p* = 1e-04; ITS: *r* = 0.05454, *p* = 1e-04). In a comparison of the Hawaiian drosophilid phylogeny and a dendrogram of Bray Curtis distance among microbiomes, the similarity between the topology of each tree is not random (16S rRNA Nye similarity = 0.3012 and ITS Nye similarity = 0.3056, when two random trees should equal 1) (**Figure 4A** and **Supp. Figure 3**). However, the degree of coevolution appears limited among fly species and microbiota based on the topological relationships of terminal taxa. The lack of phylogenetic clustering is also evident when only the most prevalent taxa are considered (**Figure 4B** and **Supp. Figure 3B**). Notably, the widely-distributed arthropod endosymbiont *Wolbachia* is present in a number of Hawaiian drosophilids. However, *Wolbachia* does not appear to be associated with speciation events among sister taxa, and instead appears clumped to certain species groups (**Figure 4C**).

**Figure 4.**
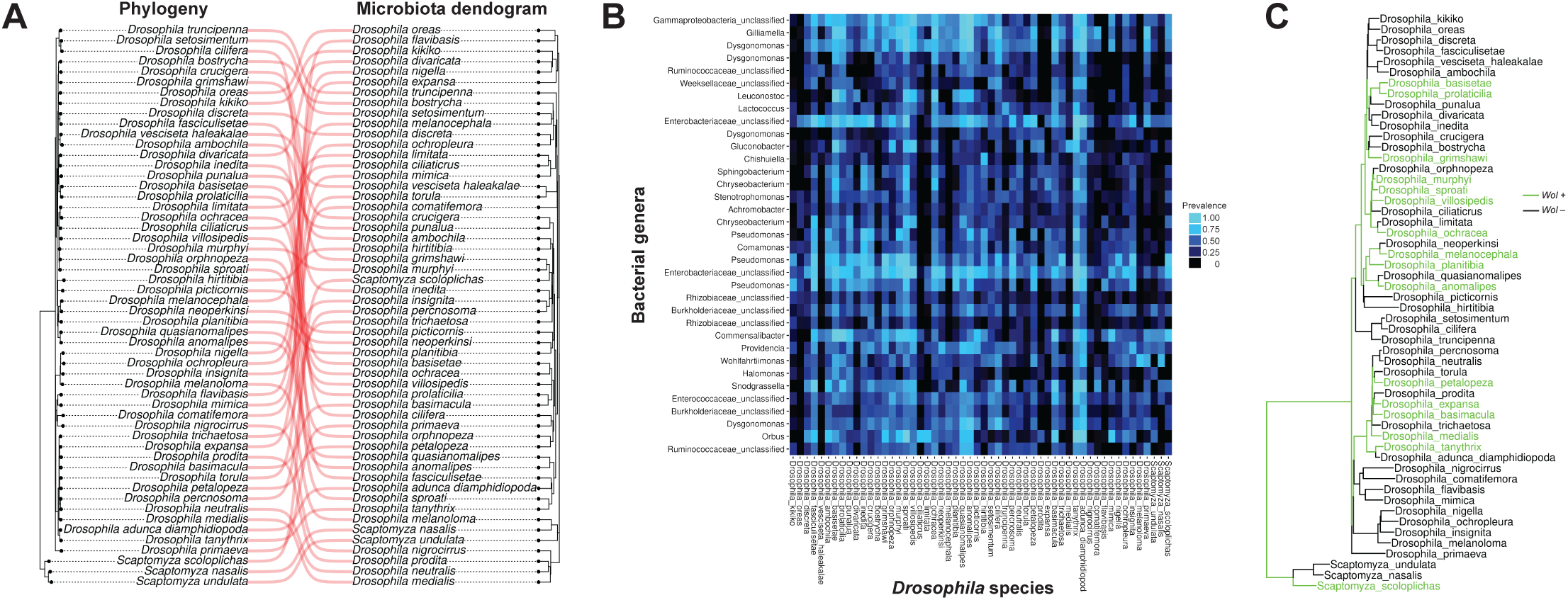
Coevolution analysis of Hawaiian *Drosophila* and microbial taxa based on 16S rRNA data. (**A**) The cophylogeny of Hawaiian *Drosophila* (left) and a dendrogram based on Bray-Curtis distance of the microbiome (right) shows weak evidence of phylosymbiosis due to the lack of topological congruence in the two trees. (**B**) The lack of a strong coevolutionary signal is also evident among the 25 most prevalent microbial taxa in Hawaiian *Drosophila*, which include a number of endosymbionts and pathogens. The fly species are listed in the same order as the fly phylogeny. (**C**) The reproductive endosymbiont *Wolbachia* is not closely associated with speciation events among sister taxa of Hawaiian *Drosophila* and appears clumped to certain species groups. The presence of *Wolbachia* was inferred for a species if *Wolbachia* was detected in any of the samples in a species.

### Community assembly of native fly gut microbiota across abiotic conditions

Given the weak effect of phylosymbiosis on microbiome composition in native flies, we employed an MRM to determine the relative contribution of genetic relatedness along with island age and other abiotic environmental conditions. Elevation (highly correlated with temperature *r* = -0.96, *p* = 2.2e-16), evapotranspiration, and island age had significant effects on structuring bacterial communities (**Table 3**, **4**; see also **Supp. Fig 2** for a visualization of each variable’s effect in a distance-based ordination). The effect of phylogenetic relatedness was non-significant, although it explained a similar amount of variation to the significant variables in the model. For fungal profiles, elevation had the strongest effect, followed by evapotranspiration, and rainfall. Interestingly, two predictor variables differed between bacteria and fungi (**Table 4**). First, island age, which had a strong effect on bacterial composition, was negligible for fungi. Second, phylogenetic relatedness was one of the strongest predictors of mycobiome composition.

**Table 3.** Multiple regression on distance matrices (MRM) analysis of only endemic Hawaiian *Drosophila* bacterial profile. ^1^Microbiome distance values among samples are based on Bray-Curtis dissimilarity values, whereas predictor variables are measured as Euclidean distance values. All predictor variables are shown and significant predictors are in bold-face (residual deviance = 8865.5).

**Table 4.** Multiple regression on distance matrices (MRM) analysis of only endemic native Hawaiian *Drosophila* fungal profiles and environmental and genetic predictors. ^1^Mycobiome distance values among samples are based on Bray-Curtis dissimilarity values, whereas predictor variables are measured as Euclidean distance values. All predictor variables are shown and significant predictors are in bold-face (residual deviance = 19919.54).

To test the possibility that bacterial communities that are specific to each site may have distinct functional properties, we compared the putative functions across significant environmental predictors. However, PICRUSt2 analysis found only slight shifts in the functional space **(Supp Figure 4)**.

### Ecological and genomic drivers of microbiome assembly

Our analysis incorporating all native fly samples provided only weak evidence for phylosymbiosis as a driver of microbial community composition. To directly assess the contribution of genetic relatedness vs. environment factors in predicting microbiome variation, we compared the microbiomes of *D. tanythrix* and *D. sproati,* two endemic species that had large sample sizes (>30 samples) and were found in 6 different sites. At least seven individuals each of *D. sproati* or *D. tanythrix* were collected from three sites on Hawai□i island: Tom’s Trail, Ola□a Pole 44, and Ka□u Forest Reserve (**Fig. 5**). Each location exhibits distinct abiotic and geographic features (**Fig. 1**). We asked first whether microbiomes of the same species differ between different sites, using PERMANOVA (**Table 5**). We observed a strong effect of site on overall microbiome variation, but not of sex. Further analysis of abiotic factors showed that only rainfall explained a significant amount of variation in the bacterial composition of *D. tanythrix* and *D. sproati* (**Table 5**). *D. sproati* from each of the field sites exhibited different bacterial and fungal profiles (**Table 6**). *D. tanythrix* showed a similar trend for 2 of 3 sites, with one exception: microbiome composition of flies from Ola□a Pole 44 and Ka□u Forest Reserve approached but did not reach significance (PERMANOVA with Bray-Curtis: 16S rRNA *p =* 0.08, ITS *p =* 0.059).

**Figure 5.**
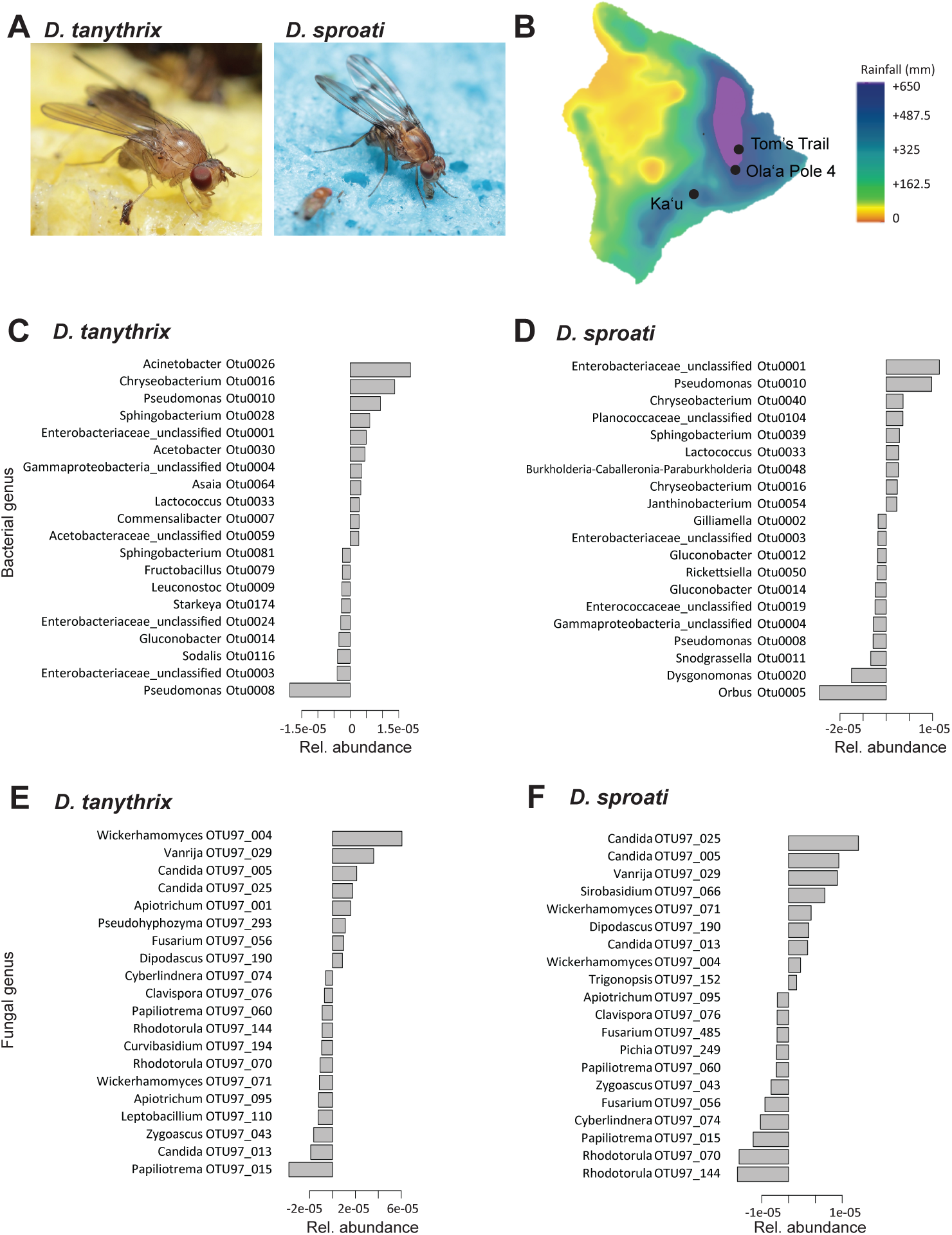
Bacterial and fungal composition are associated with rainfall levels for within-species comparisons of *D. tanythrix* and *D. sproati* microbiomes. (**A**) Images of *D. tanythrix* and *D. sproati*. (**B**) Map of Hawai‘i island showing average rainfall for January 2021 for collection sites used in analysis. Rainfall data sourced from Hawai‘i Climate Data Portal (McLean et al., 2020). (**C, D**) The top 20 bacterial taxa found in *D. tanythrix* and *D. sproati* whose abundance is associated with rainfall variation. (**E, F**) The top 20 fungal taxa whose abundance is associated with rainfall variation. Positive coefficients indicate an increase with rainfall, and negative coefficients indicate a decrease with rainfall. Photo credit: Karl Magnacca.

**Table 5.** Outcomes from PERMANOVA testing for drivers of intraspecific and intra-site variation in the bacterial microbiome of wild-caught Hawaiian *Drosophila* species.

**Table 6.** Pairwise comparisons of bacterial and fungal beta-diversity of gut microbiome for *D. sproati* and *D. tanythrix* from three locations on Hawai□i Island; T = Tom’s Trail; K = Ka□u Forest Reserve; O = Ola□a Pole 44.

Among the top bacterial taxa associated with shifts in rainfall, three OTUs common to both species were found to increase in abundance with rainfall (Otu0008: *Pseudomonas* nr. *P. alcaligenes*, Otu0014: *Gluconobacter* nr. *G. oxydans*, and Otu003: an unclassified Enterobacteriaceae) while the abundances of four species decreased with rainfall (Otu0033: *Lactococcus* nr. *L. lactis*, Otu0001: an unclassified Enterobacteriaceae, Otu0010: *Pseudomonas* nr. *P. fluorescens*, and Otu0016: *Chryseobacterium* sp.) (**Fig. 5**). For fungal profiles, evapotranspiration and rainfall were significant factors governing composition in both *D. tanythrix* and *D. sproati*, while elevation was only a significant factor for *D. sproati* (**Table 5**). Ten of the top fungal taxa were influenced by each of the three ecological variables in both fly species (OTU97_56: *Fusarium solani*, OTU97_4: *Wickerhamomyces anomalus*, OTU97_25: *Candida railenensis*, OTU97_5: *Candida asparagi*, OTU97_13: *Candida tropicalis*, OTU97_190: *Dipodascus geotrichum*, OTU97_43: *Zygoascus meyerae*, OTU97_15: *Papiliotrema flavescens*, OTU97_70: *Rhodotorula mucilaginosa*, and OTU97_74: *Cyberlindnera lachancei*) (**Fig 5; Supp. Fig 5)**.

Next, we examined whether the microbiomes of *D. sproati* and *D. tanythrix* found within the same site are similar to each other. With one exception, species from the same site tended to have similar bacterial and fungal communities (**Table 6**). No significant differences in beta-diversity were found when comparing *D. sproati* and *D. tanythrix* found within Tom’s Trail or within Ka□u Forest Reserve (16S and ITS: Bray-Curtis with PERMANOVA; p > 0.05). The only differences between *D. sproati* and *D. tanythrix* found within-site was in the bacterial profile at Ola□a Pole 44 (*p =* 0.01).

### Role of dietary habits and evidence for a core microbiome in Hawaiian *Drosophila*

For many animals, dietary specialization is facilitated through microbial symbionts that provide critical functions for nutrition acquisition or detoxification (Brune and Dietrich 2015, Haanstad and Norris 1985, Ingala et al 2021, Kwong et al 2014). Many Hawaiian *Drosophila* selectively oviposit and breed as larvae on a single native plant family (Magnacca et al 2008). To examine whether dietary specialization is associated with distinct microbial components, we compared known larval specialists with generalists. The microbiomes generally did not differ in terms of composition or bacterial or fungal taxonomic diversity (**Supp. Fig 6**) although host-plant specialists exhibited higher bacterial diversity (Shannon *p =* 0.026).

Host-plant specialization had a significant effect on both the bacterial and fungal microbiomes but only explained a small amount of among-species variation (16S data: *R*^2^ = 0.009, *p* = 0.024; ITS data: *R*^2^ = 0.007, *p* = 0.046; PERMANOVA). The assumptions of the PERMANOVA were robust since a test for homogeneity of multivariate dispersion was not significant (16S data: *p* = 0.498; ITS data: *p* = 0.79). A plot of the top 20 taxa contributing to the difference between generalists and specialists in each dataset shows a number of symbiotic taxa and fewer pathogenic bacterial taxa in the specialist species (**Figure 6**). Functional differences between generalists and specialists using PICRUSt2 suggest slight shifts in microbiome functional space (**Figure 7**).

**Figure 6.**
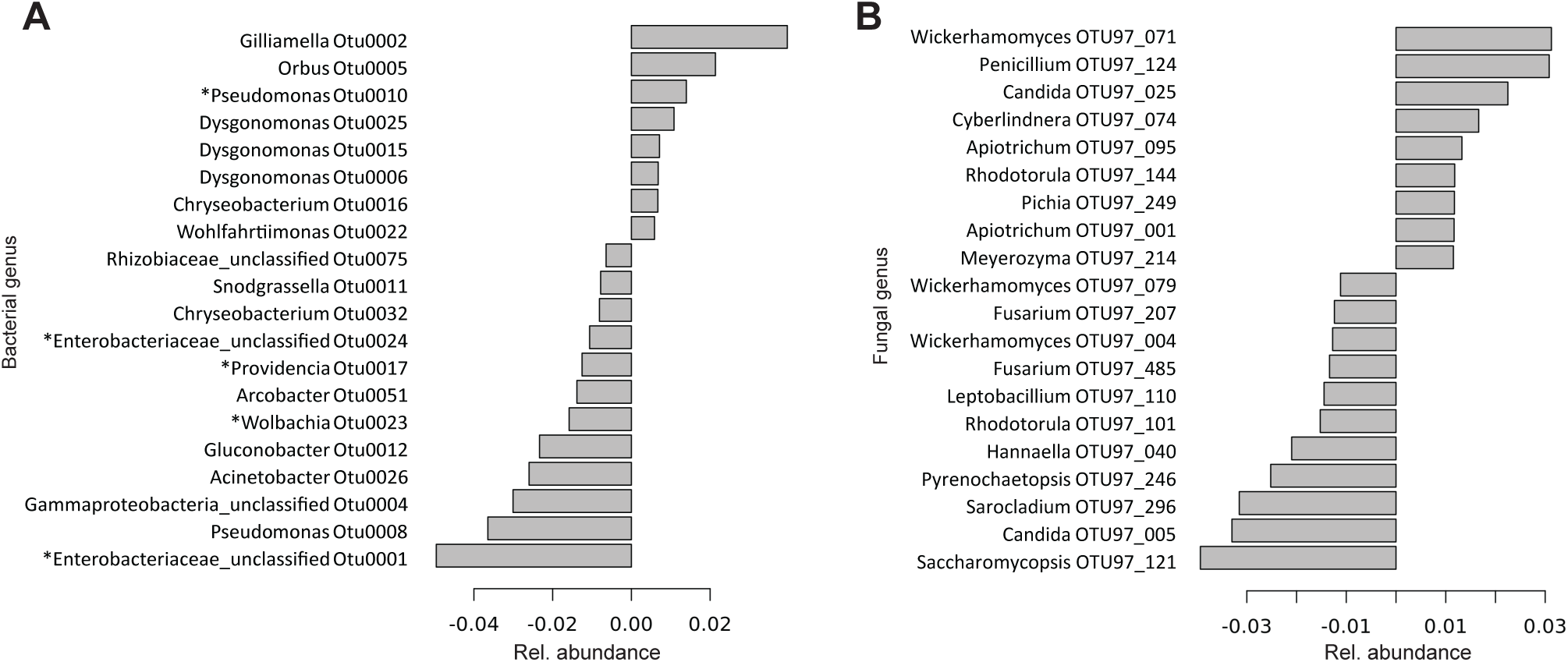
The 20 most abundant bacterial (**A**) and fungal (**B**) taxa whose abundance diverges between Hawaiian *Drosophila* dietary generalists and specialists in the picture-wing clade. Positive coefficients indicate stronger association with dietary specialists, and negative coefficients indicate a stronger association with dietary generalists; *: putative pathogens.

**Figure 7.**
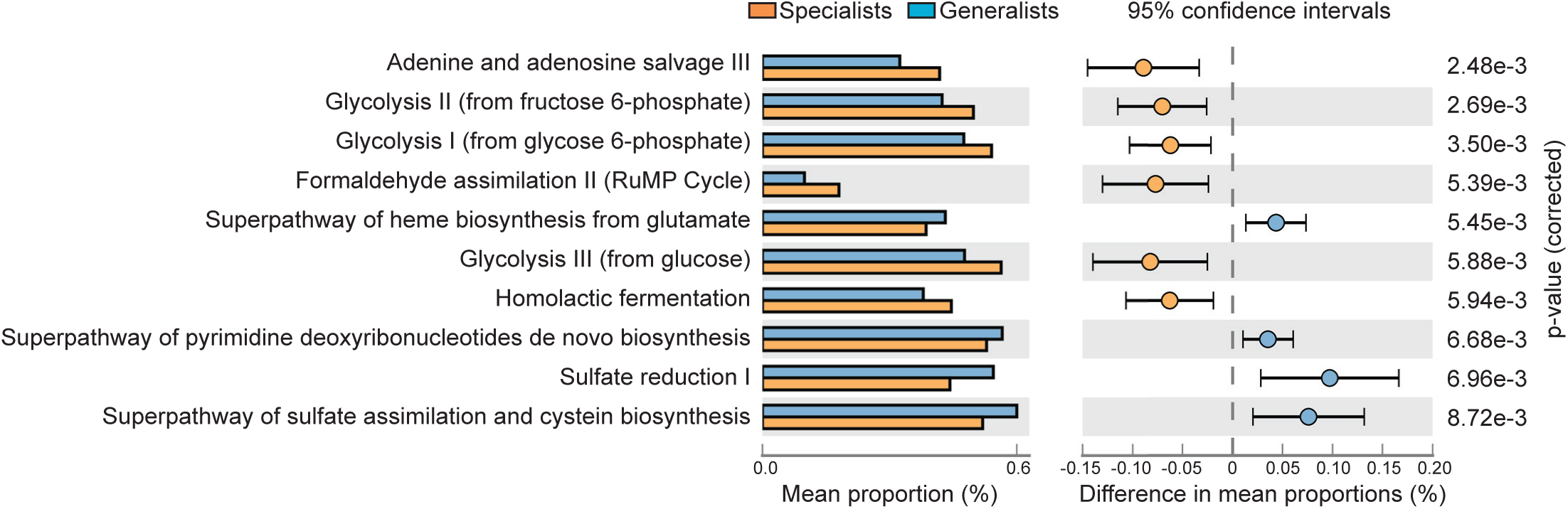
Statistically significant functional differences in specialist versus generalist Hawaiian *Drosophila* in the picture-wing clade. Functional profiles are based on KEGG annotation of microbial reference genomes and are weighted by bacterial abundance in each fly sample.

The shift in bacterial microbial diversity was associated with changes in its functions related to several notable pathways. Specialist species host microbes that are enriched for cellular energy production through central metabolic pathways, including glycolysis, homolactic fermentation, and formaldehyde assimilation II, as well as processing of nucleotides through adenine and adenosine salvage. In contrast, generalist species host microbes that are enriched for biosynthesis of heme, pyrimidine deoxyribonucleotides, and cysteine as important steps in chemical synthesis, as well as assimilation of sulfate to recover sulfur, an essential nutrient. Changes in the functional space are not due to the gain or loss of functional pathways, however, only the relative expression of pathways.

## DISCUSSION

Animals evolve in concert with the surrounding microbial world, leveraging the functions of microbial organisms to augment metabolic processes, boost immunity, and harvest nutrients from novel dietary sources. Yet, only recently have microbial factors been incorporated into models of adaptation and evolution (Henry et al 2021). The extensive radiation of the Hawaiian *Drosophila* clade provides an exemplary system to interrogate how bacterial and fungal microbiomes assemble and evolve in the context of diverse environmental conditions, dietary preferences, endemicity, and evolutionary history. Using a dataset comprised of over 500 *Drosophila* from six different Hawaiian islands, we report that abiotic factors are the main predictors of both bacterial and fungal gut microbial communities in native flies. Phylogenetic relatedness contributed to microbiome structure; however, environmental features were by far the major influencers of endemic fly microbiomes composition, consistent with previous studies of wild-caught *Drosophila* (Chandler et al 2011, Chandler et al 2012, Staubach et al 2013). By contrast, the microbiomes of cosmopolitan flies show no detectable response to changes in elevation, evapotranspiration or rainfall. Other studies have suggested that adaptation of invasive species to new niches is accompanied by the acquisition of microbes in the invaded range (Escalas et al 2022, Martignoni and Kolodny 2024, Zhang et al 2024). The invasive species in this study may have a greater resilience to variation induced by abiotic features or depend less on their microbiomes for local adaptation.

### Unique features of the Hawaiian Drosophila microbiome

Native flies exhibit different and more diverse bacterial communities compared to invasive flies but exhibit some overlap with those of other wild drosophilids, despite differences in host-plant, evolutionary lineage, and geography (Chandler et al 2011, Chandler et al 2012, Staubach et al 2013). In particular, *Enterobacteriacae*, *Providencia*, *Commensalibacter*, *Gluconobacter*, and *Dysgonomonas*, are amongst the most abundant genera found in natural populations of flies, including ones in Hawai□i. Interestingly, *Lactobacillus,* a major beneficial member of lab and wild fly microbiomes (Chandler et al 2011, Staubach et al 2013, Wong et al 2013, Wong et al 2011) was not detected in Hawaiian drosophilids. Compared to two other native arthropod radiations, *Ariamnes* spiders and true bugs, Hawaiian *Drosophila* have almost no overlap in the most abundant bacterial genera, with some exceptions such as *Pseudomonas* (Armstrong et al 2022, Poff et al 2017).

With regard to the mycobiome, *Saccahromycetes* yeast, which includes *Candida* and *Pichia* genera, were common amongst Hawaiian flies. This class of fungi is frequently associated with feeders of rotting fruit and were previously identified in a survey of cosmopolitan drosophila populations from disparate locations (Chandler et al 2012) and a study of two native Hawaiian Drosophila species (*D. neutralis*, also included in our study, and *D. imparisetae*, not in our study) (O’Connor et al 2014). The overlap in fungal features despite variation in diet, habitat, and host plants support the hypothesis that drosophilids are able to actively select for members of the *Saccahromycetes* class.

### Contribution of the microbiome to rapid adaptation

The Hawaiian *Drosophila* clade is estimated to contain as many as 1000 species at one point, originating from a single founder species (O’Grady and DeSalle 2018). To what extent could the microbiome have facilitated this explosive adaptation? Experiments with lab-raised *D. grimshawi* reveal that fecundity and mating behavior are highly dependent on microbiome composition and activity (Medeiros et al 2024). Considering the varied habitats and dietary habits exhibited by Hawaiian flies, microbes acquired from the local environment may facilitate niche expansion and local adaptation by enhancing nutrient acquisition, enabling the detoxification and nutrient harvesting of novel food sources (O’Connor et al 2014, Tefit et al 2023, Walters et al 2020). In support of this possibility, we identified slight but significant shifts in the predicted functions of bacteria from generalist and specialist flies. In particular, specialists show a shift towards pathways with greater cellular energy production, whereas generalists show a shift towards chemical synthesis. Several of the highly prevalent microbes in dietary specialist flies are also components of a core gut microbiome in insect species that have sugar-rich diets and require the degradation of complex polysaccharides or neutralization of toxic host plant allelochemicals (Zheng et al 2019). These taxa include bacteria such as *Orbus, Gilliamella, Dysgonomonas, Snodgrassella,* and *Chryseobacterium*, which are found in cactophilic *Drosophila* (Chandler et al 2011, Martinson et al 2017), as well as diverse insect species spanning Coleoptera, Hymenoptera, and Lepidoptera (Hammer et al 2020, Madden et al 2022, Shelomi et al 2023). In addition, we observed a trend towards fewer counts of bacterial pathogens in specialists. Generalist species may have higher tolerance for pathogens from encountering a wider array of host plants as larva, a pattern that has been observed for chrysomelid beetles where pathogenic bacteria dominated the microbiomes of the generalists compared to specialists (Blankenchip et al 2018).

The taxonomic diversity of fungi did not differ between endemic and invasive hosts (**Supp. Fig 1, Supp. Table 2).** Given that fungi, including yeasts, are used as a food source by *Drosophila*, it may not be surprising that similar fungal taxa are found in native and non-native flies. Fungal yeasts such as *Wickerhamomyces, Candida, Cyberlindnera*, and *Pichia* were also prevalent amongst specialists (Stefanini 2018). Several of these yeast have antimicrobial properties (Chai et al 2024), enhance *Drosophila* development (Dmitrieva et al 2019, Murgier et al 2019), or contribute to the fecundity of lab-raised Hawaiian *D. grimshawi* (Medeiros et al 2024). However, whether similar benefits are provided in the natural environment for wild populations of flies remains to be determined.

### Testing for phylosymbiosis in native Hawaiian *Drosophila*

We found weak, but significant evidence of both bacterial and fungal phylosymbiosis in native Hawaiian *Drosophila*. A pattern of phylosymbiosis has generally been supported in other animal taxa, especially mammals and birds (Harrison et al 2021) but also in the bacterial gut microbiomes of Hawaiian spiders (Perez-Lamarque et al 2022) and neotropical butterflies (Ravenscraft et al 2019). A caveat, however, is that a pattern similar to phylosymbiosis can arise from other processes, including vertical transmission of microbes through generations (Guilhot et al 2021, Guilhot et al 2023), or host filtering of microbes in its current habitat (Perez-Lamarque et al 2022).

The endosymbiont *Wolbachia* may also be implicated in the diversification of Hawaiian *Drosophila* by altering reproductive traits that could lead to reproductive isolation (Corpuz et al 2023). We found *Wolbachia* in at least some individuals of the same species identified by Corpuz et al. (2023), e.g. *D. murphyi* and *D. ochracea*, and not in others, e.g. *D. sproati* and *D. tanythrix*. As in our study, Corpuz et al. (2023) did not report finding *Wolbochia* in every specimen of a given species, supporting the idea that *Wolbochia* is not necessarily present in all individuals of a species and that modes of transmission may be both horizontal as well as vertical.

### Biogeography of Hawaiian Drosophila microbiomes

The theory of island biogeography posits that biodiversity will scale with various environmental characteristics including extent of isolation, habitat heterogeneity, and island area and age (Carey et al 2023, Gray and Cavers 2014, MacArthur and Wilson 2001). To what extent do the communities of host-associated micro-organisms follow the rules of biodiversity for macroscropic organisms? Host-associated microbial diversity has been shown to be proportional with the size of islands, both virtual, such as host plants (Chlebicki and Pawel 2007, Dinnage et al 2019, Peay et al 2007, Peay et al 2010), and actual, as shown in a study of soil from islands of the Thousand-Island Lake in China (Li et al 2020). Research on Galapagos finches revealed that island size appears to play a role in host bacterial community composition, though a number of other factors, in particular, phylosymbiosis, may explain more variation (Loo et al 2019).

We found a small but significant effect of island age on gut bacterial diversity and community makeup (**Table 1 & 3**), as well as significant differences in fungal alpha-diversity between most islands. The beta-diversity of bacterial and fungal communities differed across all five islands in our analyses. However, no clear patterns emerged associating diversity with island age or size in contrast to observations for the diversity of animals and plants (Davison et al 2018, Green et al 2004). Perhaps this outcome is not surprising considering that microbial communities are subject to host-filtering effects and have dispersal limitations (Dickey et al 2021). Differences in habitat quality and urbanization (Zhou et al 2022) also have not been accounted for in our models and could impact the diversity and composition of environmental microbiome reservoirs and, by extension, *Drosophila* microbe communities, which are largely environmentally derived (Perez-Lamarque et al 2022, Staubach et al 2013).

### Abiotic conditions are strong determinants of Hawaiian *Drosophila* microbiomes

A number of non-covarying abiotic factors were predictors of bacterial communities, particularly evapotranspiration, elevation, and island size. Similarly, evapotranspiration, elevation (highly correlated with temperature), and rainfall were predictors of fungal communities. Although little is known about the role of evapotranspiration rates and animal gut microbiomes, elevation has been found to be a predictor of gut bacterial communities in animals ranging from other *Drosophila* (Brown et al 2023) and bees (Mayr et al 2021) to fish (Bereded et al 2022). Much less is known about the role of abiotic conditions on the gut mycobiome though abiotic conditions likely play a role in community assembly, as shown for mesquite spiny lizards (Montoya-Ciriaco et al 2020). When comparing only *D. sproati* and *D. tanythrix*, two species we collected from multiple locations on Hawai□i Island, rainfall was a strong predictor of bacterial community diversity, despite rainfall not being a predictor of bacterial composition for our larger dataset with all species included. *D. tanythrix* and *D. sproati* diverged between 7.5 - 11 Mya (Church and Extavour 2022, Magnacca and Price 2015). Additionally, each species has access to distinct microbial reservoirs: *D. tanythrix* utilize leaves of the host plant *Cheirodendron trigynum* (□Ōlapa) on the ground for breeding, whereas *D. sproati* lay eggs in decaying *C. trigynum* bark and females likely do not approach leaf litter (Magnacca et al 2008). Despite these differences in evolutionary history and lifestyle, the microbiomes of *D. sproati* and *D. tanythrix* found within a site were not significantly different. By contrast, microbial communities were distinct for populations of the same species found in different sites. These observations lend additional support for the hypothesis that abiotic conditions, rather than host species identity, are the main driver of microbiome community assembly in native Hawaiian *Drosophila*.

## CONCLUSION

The Hawaiian *Drosophila* clade is an iconic model for understanding how unique features of island ecology such as diverse habitats, steep environmental gradients, and geographic isolation contribute to rapid adaptation and speciation. Along with these features, the microbial environment is likely to play a prominent role in shaping evolution. Here, our microbiome survey of native and non-native flies from across the Hawaiian archipelago reveal that the microbiomes of endemic Hawaiian *Drosophila* are distinct from those of non-native species in terms of composition, greater diversity, and response to abiotic conditions. We found only a weak signature of phylosymbiosis in the gut microbial communities of native *Drosophila*. In contrast, elevation, evapotranspiration, and rainfall appear to be more influential as drivers of bacterial and fungal species in endemic, but not invasive, drosophilids. Microbiomes of the same species differ between sites on the same island, illustrating the robust influence of environmental features. The concurrent evolution of gut microbial communities, along with their hosts, has been hypothesized to play a role in local adaptation and even ecological speciation (Rennison et al 2019). The presence of local, environmentally-specific microbes may facilitate adaptation by enhancing host ability to exploit local food sources, enhancing fecundity, altering metabolism, or influencing mating behavior (Medeiros et al 2024). Future functional studies comparing the microbiomes and behaviors of wild caught species found on different islands would allow this hypothesis to be tested. Additionally, careful characterization of host plant microbes will help to clarify the extent to which host plants serve as microbial reservoirs for Hawaiian flies and contribute to their diversification (Ort et al 2012).

## ACKNOWLEDGEMENTS

We are grateful to the following individuals for assisting with field work: Keahi Bustamente, Will Haines, Dennis Hokama (Lāna’i lodging), Kevin & Linda Jenkins (Maui lodging), Kelli Konicek, Karl Magnacca, David Medeiros, Ann Nguyen, and Michelle Smith. We thank Kelli Konicek, Karl Magnacca, and Ken Kaneshiro for helpful discussions; Valerio Stacconi for use of the *D. suzukii* image; the Hawai‘i Invertebrate Program and Joel Sartore and the Photo Ark project for use of the *Drosophila grimshawi* image. Additionally, we thank the following agencies for permitting us to conduct this research: Division of Forestry and Wildlife (State of Hawai□i and local offices on Kaua□i, O□ahu, Lānai, and Hawai□i Island), TNC Molokai, East Maui Irrigation, Hawai□i Volcanoes National Park, Hawai□i Experimental Tropical Forest, TNC Maui, Koke□e State Park, and Pūlama Lāna□i. This research was conducted in compliance with the State of Hawai‘i Division of Forestry and Wildlife regulations under the authority of Native Invertebrate Research Permits I302, I2456, I2674, and 12978. Access to field sites was authorized under KPI-2019-214 (Kauai 2019), KPI2021-245 (Kauai 2021), K2019-4061cc (Koke’e SP 2019), I2456 (Big Island 2019), HAVO-2021-SCI-0013 (Hawai‘i Volcanoes National Park). Samples for high throughput sequencing were processed by the UHM Microbial Genomics and Analytical Laboratory core (supported by NIH NIGMS P20GM125508 and P20GM139753). This work was funded by the National Science Foundation Grant No. 2025669 (MJM, JYY), National Institutes of Health Grant No. P20GM125508 (MJM, JYY), Hawai□i Community Foundation Grant No. 19CON-95452 (MJM, JYY).

## AUTHOR CONTRIBUTIONS

MJM, SDS, DKP, JYY designed and performed research, analyzed data, and contributed to the writing and editing of the paper.

## DATA AVAILABILITY STATEMENT

Sequence data will be submitted to the NCBI Sequence Read Archive. Data for Hawai‘i climate variables and topographic data were sourced from the Hawai‘i Climate Data Portal (McLean et al., 2020), TessaDEM database (https://tessadem.com/) and OpenTopography (USGS 2018).

## Supplemental figure legends

**Supp. Fig. 1.** Alpha diversity analyses of bacteria and fungi found in Hawaiian *Drosophila* (HD) and non-native *Drosophila* (nonHD). (**A**) Hawaiian *Drosophila* (n=440) exhibit higher bacterial taxonomic richness and evenness compared to non-native flies (n=51). (**B**) Fungal communities of Hawaiian *Drosophila* (n=453) have slightly higher taxonomic richness compared to non-native flies (n=57) but similar evenness; *ns*: not significant.

**Supp. Fig. 2.** Ordinations employing a distance-based redundancy analysis (dbRDA) of Bray-Curtis distance of Hawaiian *Drosophila* bacterial communities (left **A-C**, 16S rRNA dataset; right **D-F**, ITS dataset) in relation to ecological predictors. For endemic Hawaiian species, (**A, D**) elevation, (**B, E**) evapotranspiration, (**C**) island age (for bacteria) and (**F**) rainfall (for fungi) had significant effects on microbiome composition. For non-native species (A’ – F’), elevation, evapotranspiration, island age, and rainfall were not significant predictors of microbiome variation. Rainfall had an effect on non-native mycobiomes but this outcome is likely driven by an outlier.

**Supp. Fig. 3.** Coevolution analysis of Hawaiian *Drosophila* and fungal taxa based on ITS data. (**A**) The phylogeny of Hawaiian *Drosophila* (left) and a dendogram based on Bray-Curtis distance (right) shows weak evidence of phylosymbiosis as evidenced by the topological switching shown by the red lines. (**B**) Examination of the three most prevalent fungal taxa in Hawaiian *Drosophila* reveals a lack of a strong coevolutionary signal.

**Supp. Fig. 4.** Principal component analyses examining the relationship between Hawaiian *Drosophila* bacterial microbiome function and ecological predictors. The variables (A) elevation, (B) evapotranspiration, and (C) island age show slight divergence in functional microbiome composition with each variable.

**Supp. Fig. 5.** Fungal composition is associated with abiotic factors for within-species comparisons of *D. tanythrix* and *D. sproati* microbiomes. (**A**) Map of Hawai‘i Island showing the average annual evapotranspiration levels for 2014. Data sourced from Hawai‘i Climate Data Portal (McLean et al., 2020). (**B, C**) The top 20 fungal taxa found in *D. tanythrix* and *D. sproati* whose abundance is associated with evapotranspiration. (**D**) Topographic map of Hawai‘i Island showing elevation at collection sites. Data sourced from TessaDEM database (<u>https://tessadem.com/</u>) and OpenTopography (USGS 2018). (**E**) The top 20 fungal taxa found in *D. sproati* whose abundance is associated with elevation.

**Supp. Fig. 6.** Comparison of microbiome communities from generalist and specialist Hawaiian *Drosophila*. (**A**) Alpha diversity analyses comparing the 16S rRNA profiles of generalist (n=30) and specialist flies (n=184). Only Shannon diversity differed significantly (Mann-Whitney test). (**B**) Non-multidimensional scaling (NMDS; based on Bray-Curtis distances) indicate no significant differences in beta diversity. Ellipses represent significance at the 0.05 confidence interval. (**C, D**) Alpha and beta diversity analyses comparing the ITS profiles of generalist (n=42) and specialist flies (n=190). No significant differences were found in terms of taxonomic diversity or composition.

## Supplemental table legends

**Supp. Table 1**. Summary of collection site locations and species collected at each site.

^1^PWD: picture-wing *Drosophila*

^2^Based on CO1 barcoding.

^3^Number of individuals used for amplicon sequencing.

**Supp. Table 2**. Prevalent microbial taxa found in endemic Hawaiian *Drosophila* species.

**Supp. Table 3.** Multiple regression on distance matrices (MRM) analysis of only non-native Hawaiian *Drosophila* bacterial profiles.

^1^Microbiome distance values among samples are based on Bray-Curtis dissimilarity values, whereas predictor variables are measured as Euclidean distance values. All predictor variables are shown (residual deviance = 8865.5).

