## Supplementary figures and images for "Abiotic factors are the primary determinants of endemic Hawaiian *Drosophila* microbiome assembly"

### Supp. Fig. 1

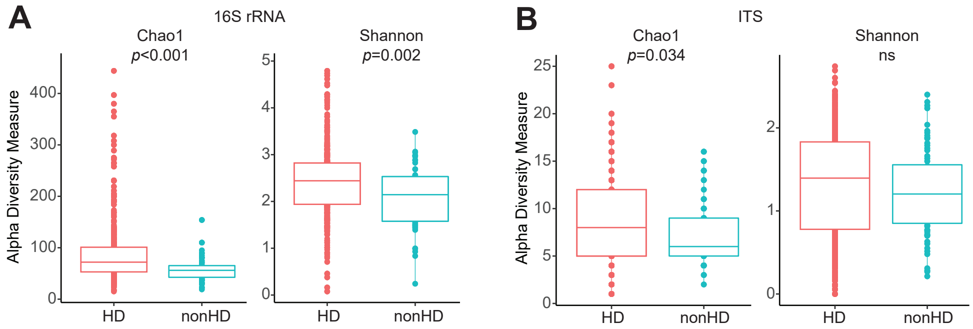

### Supp. Fig. 2

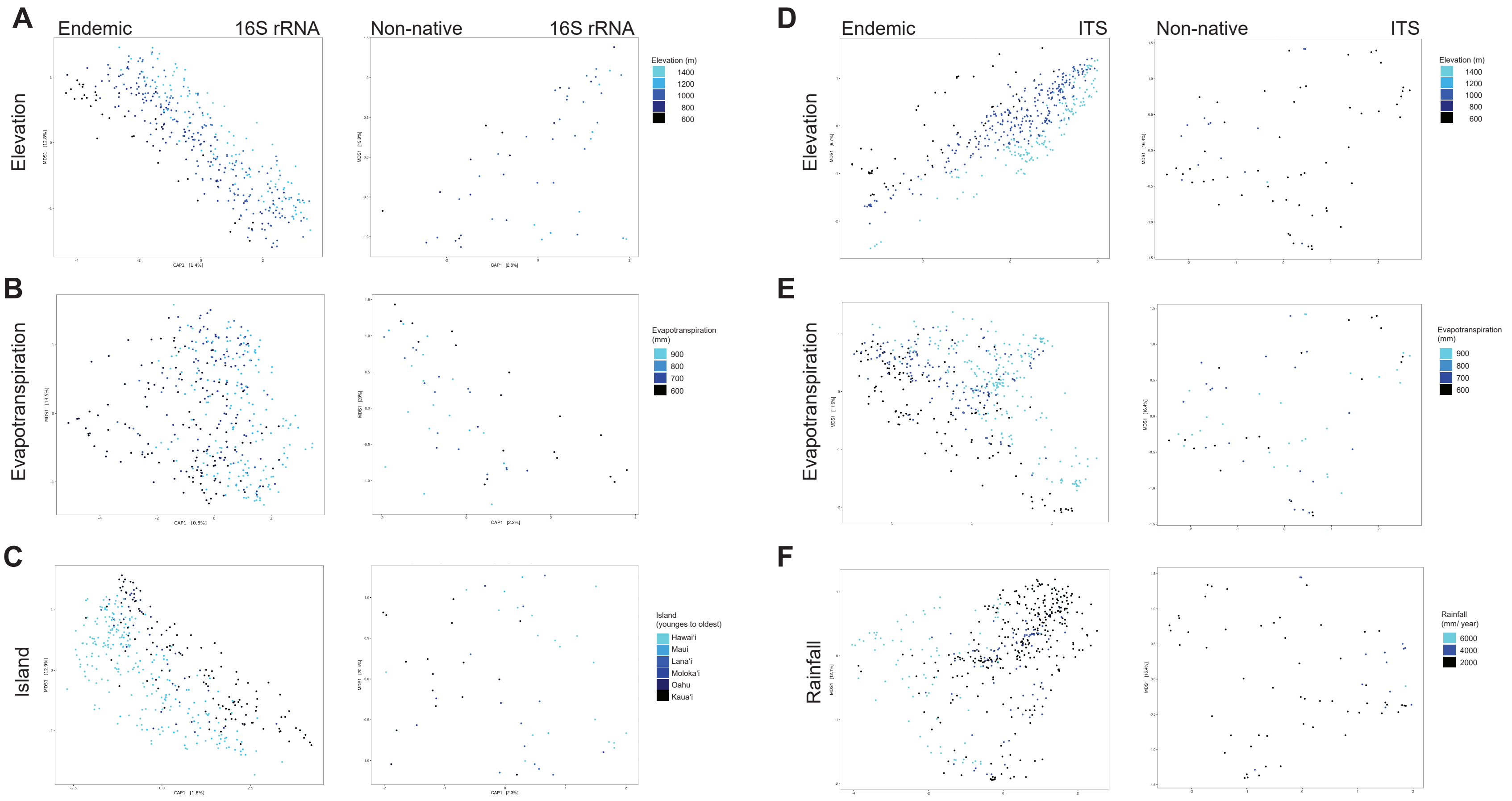

### Supp. Fig. 3

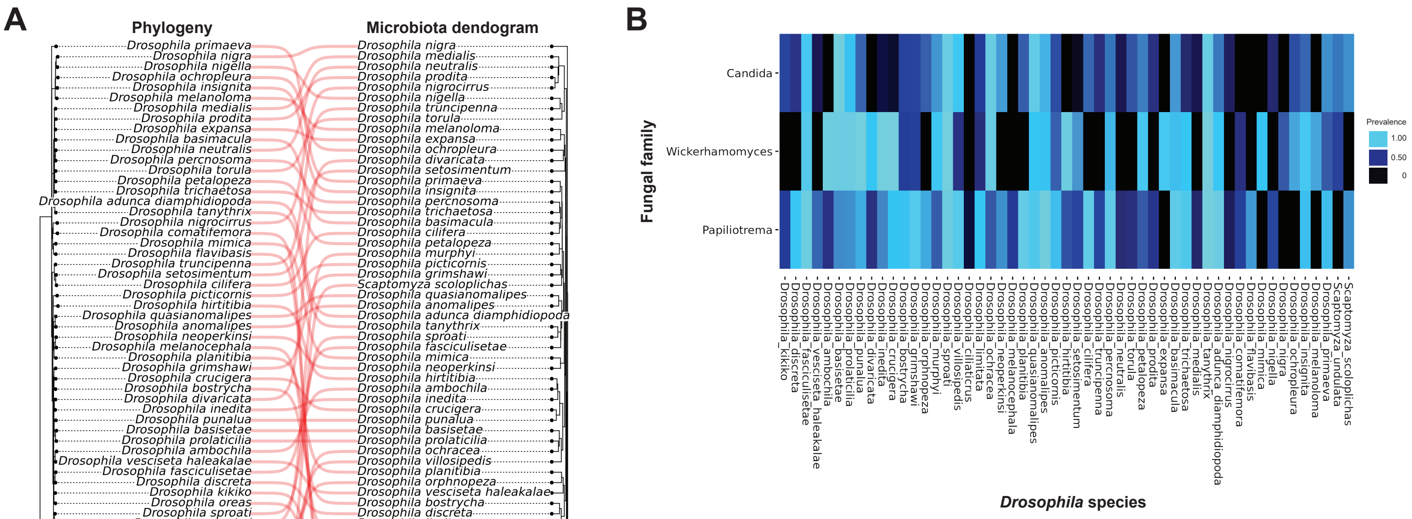

### Supp. Fig. 4

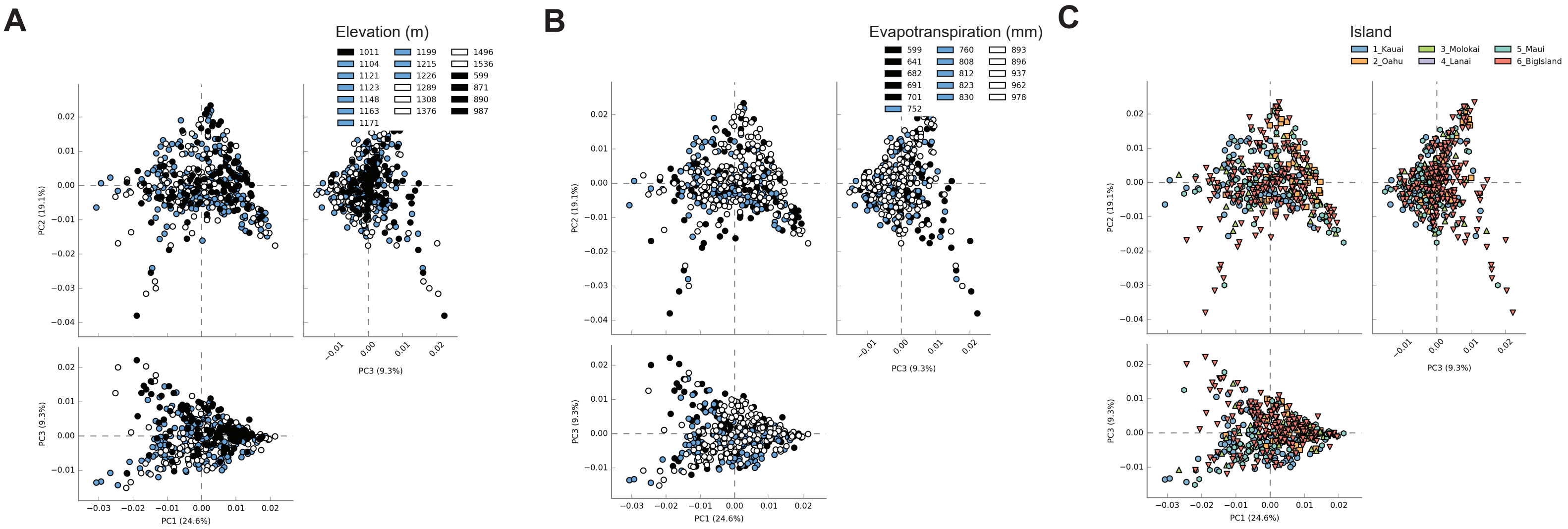

### Supp. Fig. 5

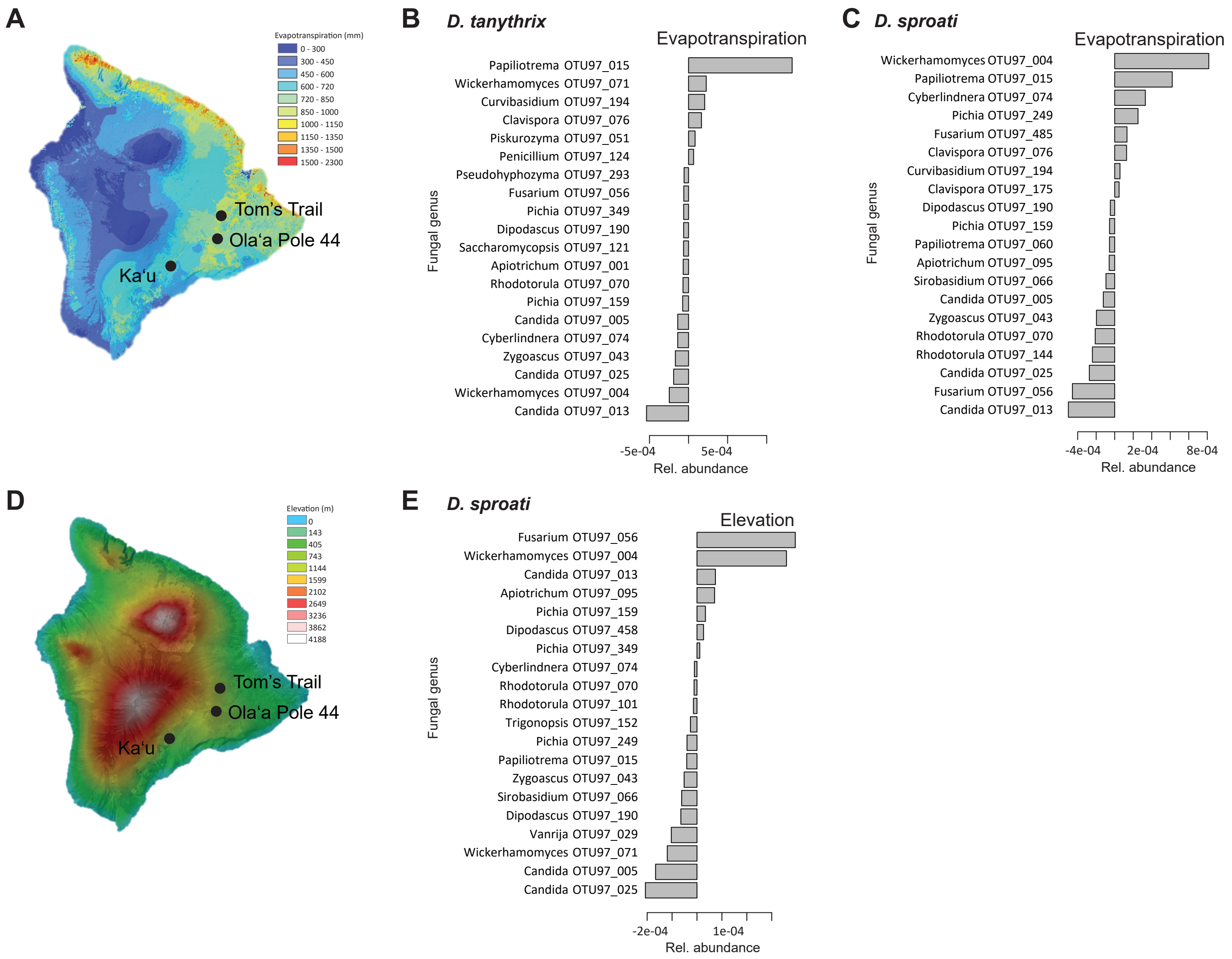

### Supp. Fig. 6

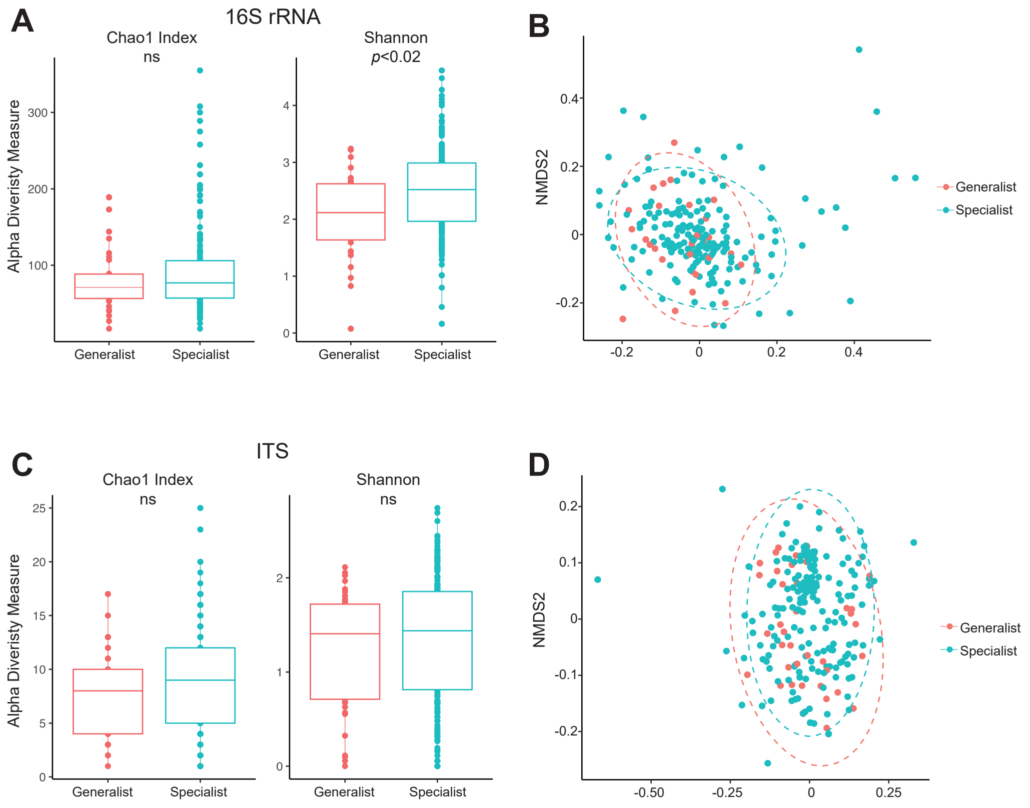
