## Supplementary material for "Abiotic factors are the primary determinants of endemic Hawaiian *Drosophila* microbiome assembly": Supp. Table 3

**Supp. Table 3.** Multiple regression on distance matrices (MRM) analysis of only non-native Hawaiian *Drosophila* bacterial profiles.

| **Variable^1^** | **β-value^2^** | ***p*-value** |
| --- | --- | --- |
| Intercept | 1.459 | 0.855 |
| Sex | -0.010 | 0.856 |
| Elevation | 0.126 | 0.178 |
| Rainfall | -0.003 | 0.977 |
| Evapotranspiration | -0.035 | 0.561 |
| Island Age | 0.006 | 0.919 |

^1^Microbiome distance values among samples are based on Bray-Curtis dissimilarity values, whereas predictor variables are measured as Euclidean distance values. All predictor variables are shown (residual deviance = 8865.5).
